# The proteomic and transcriptomic landscapes altered by Rgg2/3 activity in *Streptococcus pyogenes*

**DOI:** 10.1101/2022.05.06.490990

**Authors:** Britta E. Rued, Caleb M. Anderson, Michael J. Federle

## Abstract

*Streptococcus pyogenes*, otherwise known as Group A Streptococcus (GAS), is an important and highly adaptable human pathogen with the ability to cause both superficial and severe diseases. Understanding how *S. pyogenes* senses and responds to its environment will likely aid in determining how it causes a breadth of diseases. One regulatory network involved in GAS’s ability to sense and respond to the changing environment is the Rgg2/3 quorum sensing (QS) system, which responds to metal and carbohydrate availability and regulates changes to the bacterial surface. To better understand the impact of Rgg2/3 QS on *S. pyogenes* physiology, we performed RNA-seq and TMT-LC-MS/MS analysis on cells in which this system was induced or disrupted. Primary findings confirmed that pheromone stimulation in wildtype cultures is limited to the induction of operons whose promoters contain previously determined Rgg2/3 binding sequences. However, supplementing exogenous pheromone to a deletion mutant of *rgg3*, a strain that endogenously produces elevated amounts of pheromone, led to extended alterations of the transcriptome and proteome, ostensibly by stress-induced pathways. Under such exaggerated pheromone conditions (Δ*rgg3*+SHP), a connection was identified between Rgg2/3 and the stringent response. Mutation of *relA*, the bifunctional guanosine tetra- and penta-phosphate nucleoside synthetase/hydrolase, and alarmone synthase genes *sasA* and *sasB*, impacted culture doubling times and disabled induction of Rgg2/3 in response to mannose, while manipulation of Rgg2/3 signaling modestly altered nucleotide levels. Our findings indicate that excessive pheromone production or exposure places stress on GAS resulting in an indirect altered proteome and transcriptome beyond primary pheromone signaling.

**IMPORTANCE:** *Streptococcus pyogenes* causes several important human diseases. This study evaluates how the induction or disruption of a cell-cell communication system alters *S. pyogenes*’s gene expression and, in extreme conditions, its physiology. Using transcriptomic and proteomic approaches, the results define the pheromone-dependent regulon of the Rgg2/3 quorum sensing system. In addition, we find that excessive pheromone stimulation, generated by genetic disruption of the system, leads to stress responses that are associated with the stringent response. Disruption of this stress response affects the ability of the cell-cell communication system to respond under certain conditions. These findings assist in the determination of how *S. pyogenes* is impacted by and responds to non-traditional sources of stress.

## INTRODUCTION

*Streptococcus pyogenes*, otherwise known as Group A Streptococcus (GAS), causes millions of infections per year worldwide (1, 2), resulting in an array of disease states, from superficial tonsilitis to severe infections such as necrotizing fasciitis (3, 4). Genetic programs that regulate virulence attributes of GAS are responsive to the microorganism’s environment, and its metabolic status, through intercellular signaling networks. Quorum sensing (QS), a form of intercellular communication, is used by GAS to regulate gene expression in response to molecular signals produced by bacteria (5–11). QS activity is often coupled to the status of the environment, where characteristics such as nutrient availability or pH can impact expression or action of QS-centric genes. A conserved network of QS pathways in GAS involves Rgg transcription factors, which are receptors for peptide pheromones. Rgg proteins are a part of a larger family of pheromone-responsive proteins known as RRNPP (Rap, Rgg, NprR, PlcR, PrgX) (12). GAS possesses four Rgg proteins, RopB (Rgg1), Rgg2 and Rgg3 (Rgg2/3), and ComR (Rgg4) (13). Several studies have documented their roles in virulence, biofilm formation, and lysozyme resistance (5, 6, 8, 9, 11, 14, 15).

Rgg2 and Rgg3 are allosteric regulators of transcription that respond indiscriminately to the short hydrophobic peptides SHP2 and SHP3. The active forms of the peptides differ by only one conserved amino acid (SHP2: DIIIIVGG; SHP3: DI<u>L</u>IIVGG), and empirical evaluation has determined they have equivalent inducer activities; for simplicity we refer to them as SHP (5, 9, 14, 16–21). Induction of gene transcription requires Rgg2, which when bound to SHP, acts as a transcriptional activator. Deletion of *rgg2* (Δ*rgg2*) disables transcription induction, where target genes remain silent or express only at very low levels (16). In contrast, Rgg3 represses transcription until it binds to SHP, whereupon repression is abrogated (16, 17). Environmental signals also integrate into the Rgg2/3 circuit. Depleting growth media of manganese or iron, or provision of mannose as a primary carbon source, each have the effect of inducing QS signaling (14). We have suggested these conditions resemble the nutrient status seen in the host where GAS communication may enhance fitness and adaptability.

Two polycistronic operons are known regulatory targets of Rgg2/3, defined in part by an Rgg2/3 binding sequence that overlaps a typical -35 promoter element (17). At one locus, *shp2* is followed by *spy49*_*0414c* (*stcA*). The other locus encodes *shp3*, nine putative enzymes, and a major facilitator superfamily (MFS) transporter (*spy49*_*0450* – *spy49_0460)* (Chang *et. al*, 2022, submitted). *stcA* is associated with enhanced lysozyme resistance and biofilm formation, whereas *spy49_0450-0460* is required for QS-induced immune suppression (5, 9, 14).

Given the complex nature of identified phenotypes associated with this QS system, we wondered if other genes are under SHP and Rgg2/3 control. To assess this, we performed transcriptomic and proteomic analyses to observe responses of wildtype NZ131 and isogenic Δ*rgg3* and Δ*rgg2* mutants in the presence and absence of SHP. We find that the transcriptomic and proteomic responses of wildtype stimulated by SHP are restricted to genes previously associated with the system. However, a mutant of *rgg3*, which produces elevated amounts of pheromone due to its de-repressed state, when further stimulated with exogenous SHP, results in differential gene and protein expression extending beyond patterns observed for stimulated wildtype, likely as an indirect stress response.

## RESULTS

### RNA-seq identifies the core regulon for Rgg2/3 quorum sensing

Given that multiple environmental inputs impact Rgg2/3 signaling, and because phenotypes associated with this QS system are complex (e.g., immunosuppression, lysozyme resistance) and likely to involve several genes, we evaluated gene and protein expression patterns when GAS NZ131 (serotype M49) was induced by SHP and in isogenic mutant backgrounds containing Δ*rgg2* or Δ*rgg3*. Direct sequencing and quantitation of genomic mRNA transcripts with analysis to identify differentially expressed genes (DEGs) between unstimulated cultures and those treated with SHP led to the determination that 11 genes were induced >240-fold and constitute the core Rgg2/3 regulon (Fig. 1A-B, Table 1, Data Set S1). Confirming prior reports (5, 9, 16), *spy49_0414c* (*stcA*) and *spy49_0450-0460* were expressed from transcription start sites directly upstream of either *shp2* or *shp3* and appear to be co-expressed as polycistronic operons. Three additional genes were significantly differentially regulated: *spy49_0461c* and *0462c* are hypothetical genes encoded immediately downstream and on the opposite strand from the *0450-0460* operon, and *spy49_0677c* (transposase, *ISS1mu1*) was found to be differentially expressed ∼4-fold and barely meeting statistical criteria. A similar list of genes was identified when comparing Δ*rgg2* to Δ*rgg3* strains, which are effectively non-inducible and auto-inducible strains, respectively (Fig. 1C-D, Data Set S1). Aside from this, principal component analysis indicates that Δ*rgg3* strongly mimics WT induced with SHP, whereas the Δ*rgg2* strain replicates WT grown without SHP induction in CDM medium (Fig. 3A). These observations reinforce our previous suggestion that Δ*rgg3* exists in a QS-ON status, where Δ*rgg2* remains in a QS-OFF state.

**FIGURE 1:**
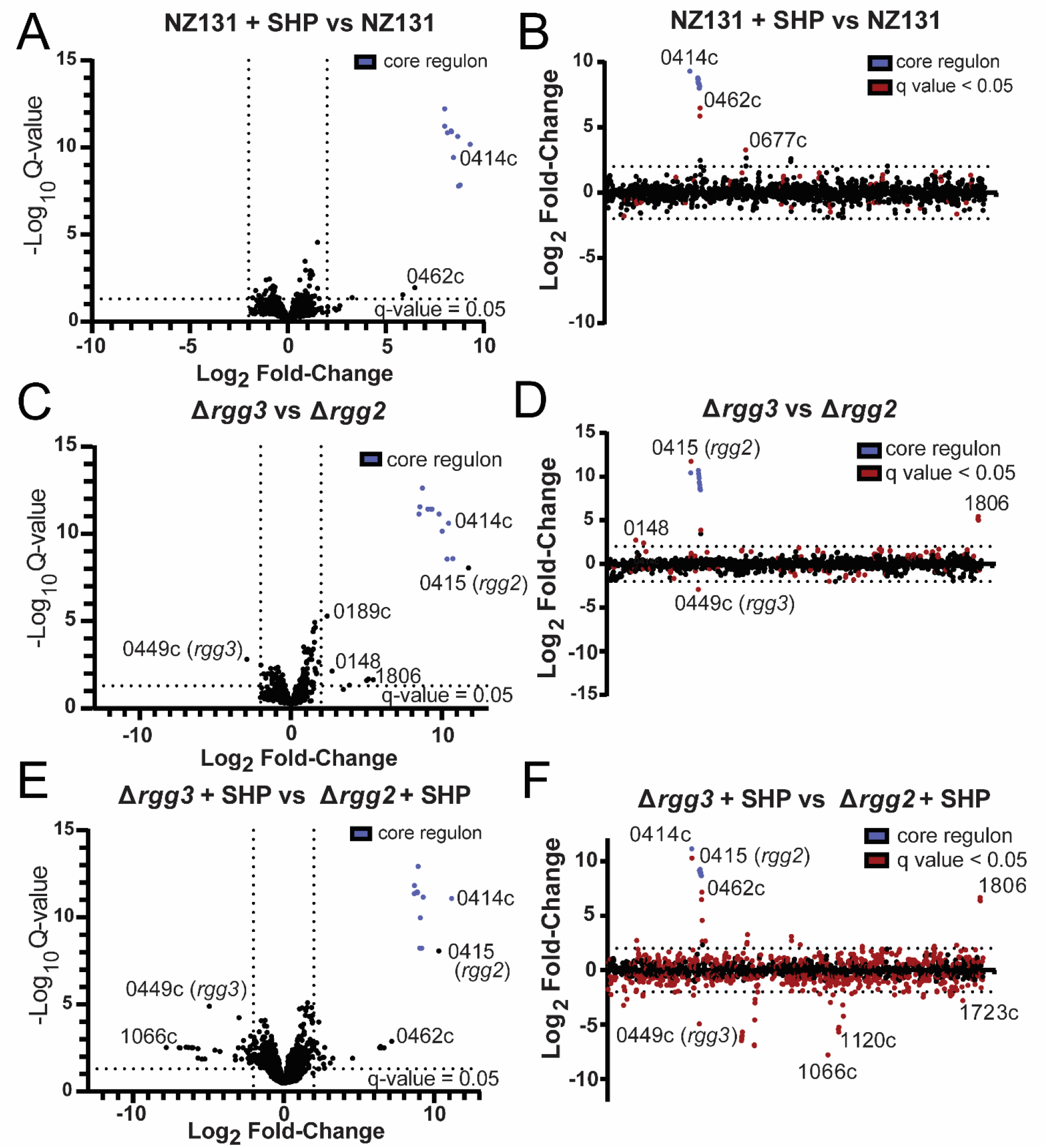
Transcriptomic changes in response to pheromone stimulation. For a complete list of differentially regulated genes and RNA-seq data, see Data Set S1. Volcano plots (panels A, C, E) indicate transcript fold-changes and *q*-value significance. The core Rgg2/3 regulon genes are indicted by blue points. Dotted lines indicate threshold values for log2 fold-change of transcript counts (minimum 4-fold) or log10 q-values (<0.05) (Panels B, D, F). Log2-fold changes of transcript counts plotted as a function of the NZ131 linear genome map. Red dots indicate significant q-values <0.05. Dotted line indicates threshold > 4-fold differential transcript counts. A-B) NZ131 vs. NZ131 with supplemented 100 nM SHP. (C-D) Δ*rgg3* vs Δ*rgg2.* (E-F) Δ*rgg3* vs. Δ*rgg2* cultures, each stimulated with 100 nM SHP.

**FIGURE 2:**
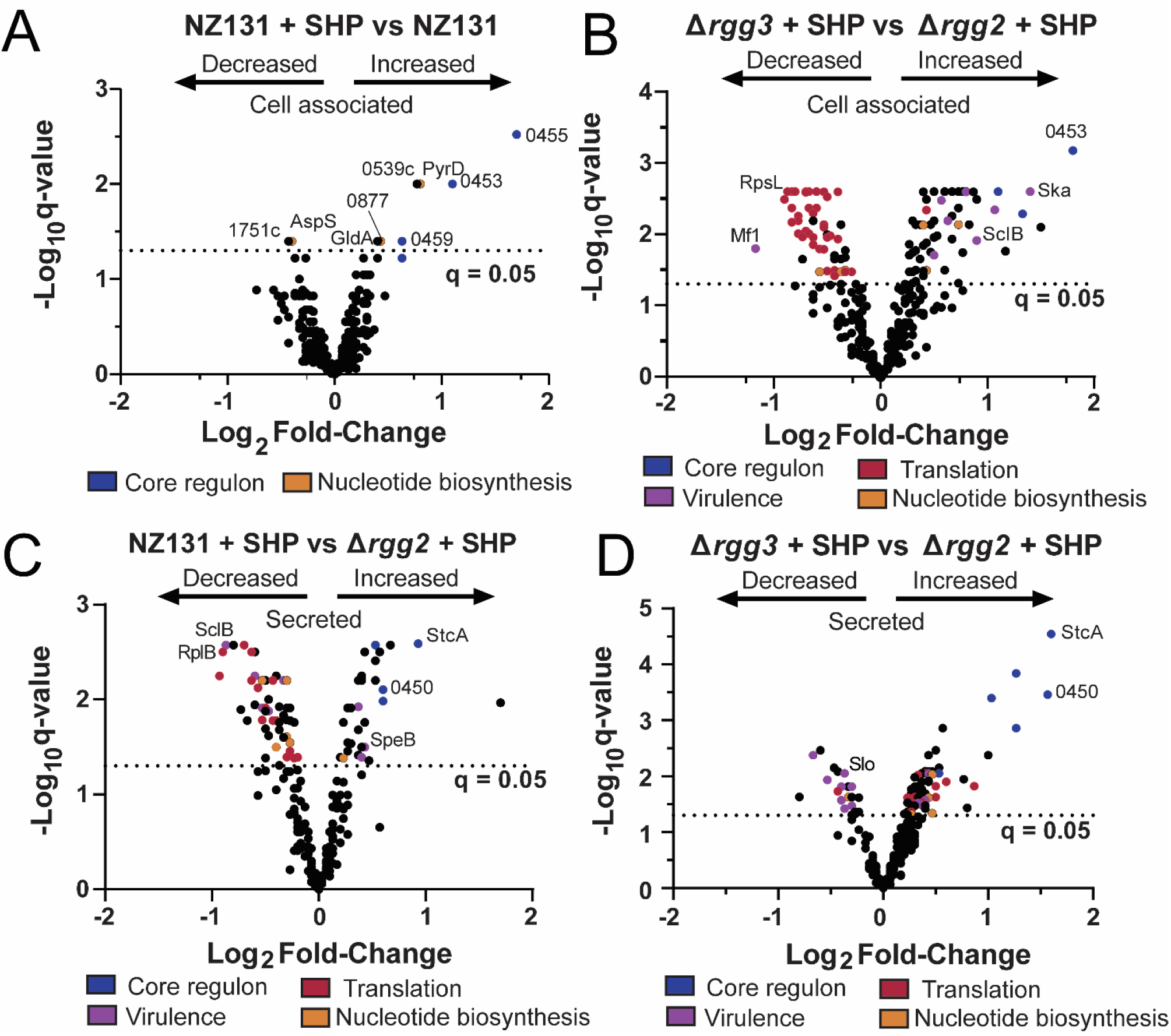
Volcano plots of proteomics results. The legends below the graphs indicate processes or operons in which proteins are involved. Blue, core Rgg2/3 regulon; purple, virulence; red, translation; orange, nucleotide biosynthesis. Proteins of particular interest are also indicated on each graph. For a complete list of differentially expressed proteins and TMT-MS/MS results, see Data Sets S2 and S3. A) Differentially expressed cell-associated proteins in wild-type NZ131 + SHP vs NZ131. B) Differentially expressed cell-associated proteins in Δ*rgg3* + SHP vs Δ*rgg2* + SHP. C) Differentially expressed secreted proteins in wild-type NZ131 + SHP vs Δ*rgg2* + SHP. D) Differentially expressed secreted proteins in Δ*rgg3* + SHP vs Δ*rgg2* + SHP.

**FIGURE 3:**
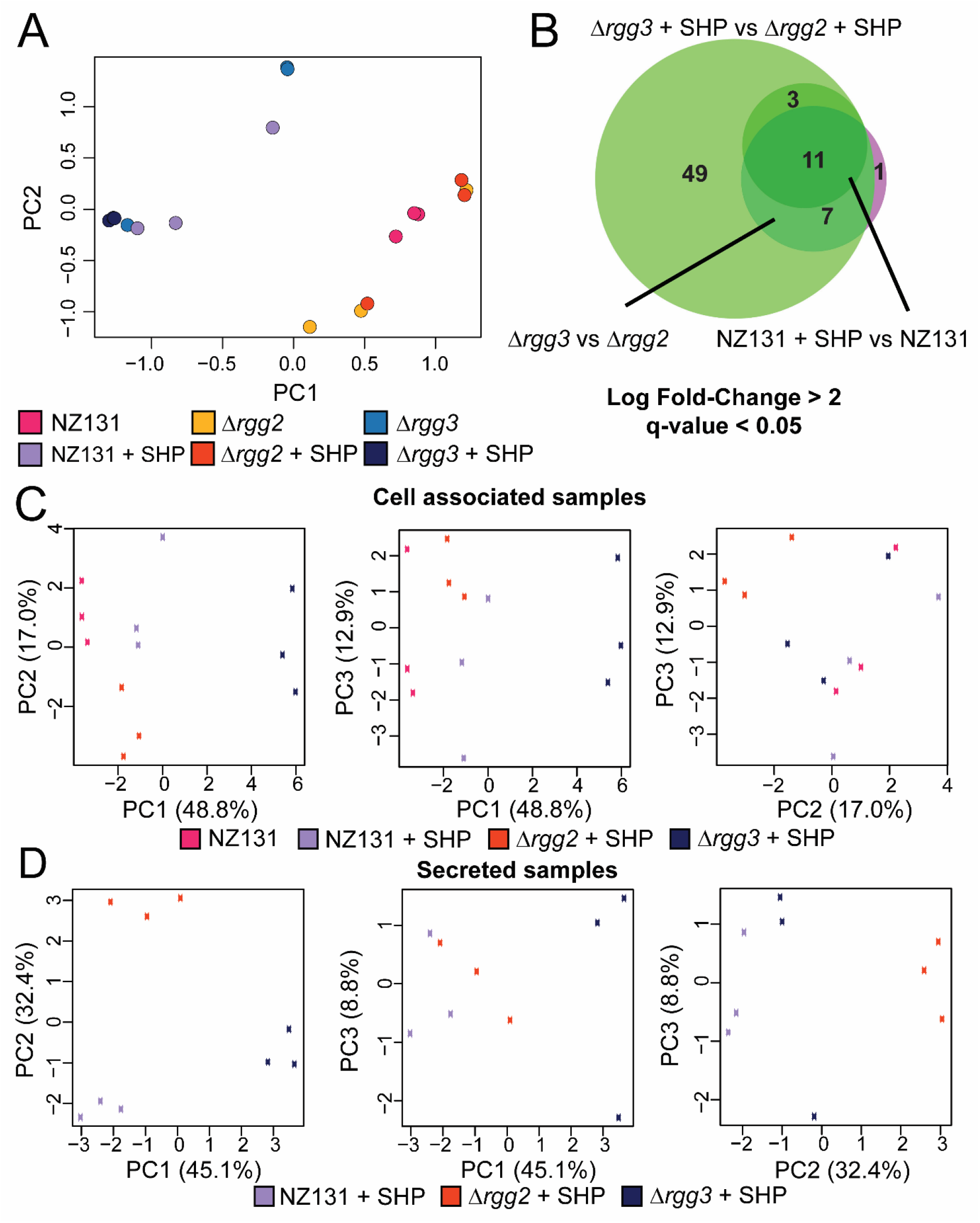
PCA analysis of RNA-seq and proteomics results, and additional RNA-seq data. For a complete list of differentially expressed transcripts and proteins, see Data Sets S1-S3. A) Principle component analysis of RNA-seq results. The legend below the graph indicates the color of each sample, for which three biological replicates were harvested. B) Venn diagram of RNA-seq results, demonstrating a core regulon of 11 genes for the Rgg2/3 system. NZ131 + SHP vs NZ131 is indicated by the darker green circle, Δ*rgg3* vs Δ*rgg2* by the pink-purple circle, and Δ*rgg3* + SHP vs Δ*rgg2* + SHP by the light green circle. Log fold-change greater than 2 and q-value of less than 0.05 was used as a cutoff for significance. 11 genes are found to be differentially regulated in all conditions. C) Three-way principal component analysis of cell associated samples proteomics results. The legend below the graphs indicates the corresponding sample with the box color on each graph. D) Three-way principal component analysis of secreted samples proteomics results. The legend below the graphs indicates the corresponding sample with the box color on each graph.

**TABLE 1.**
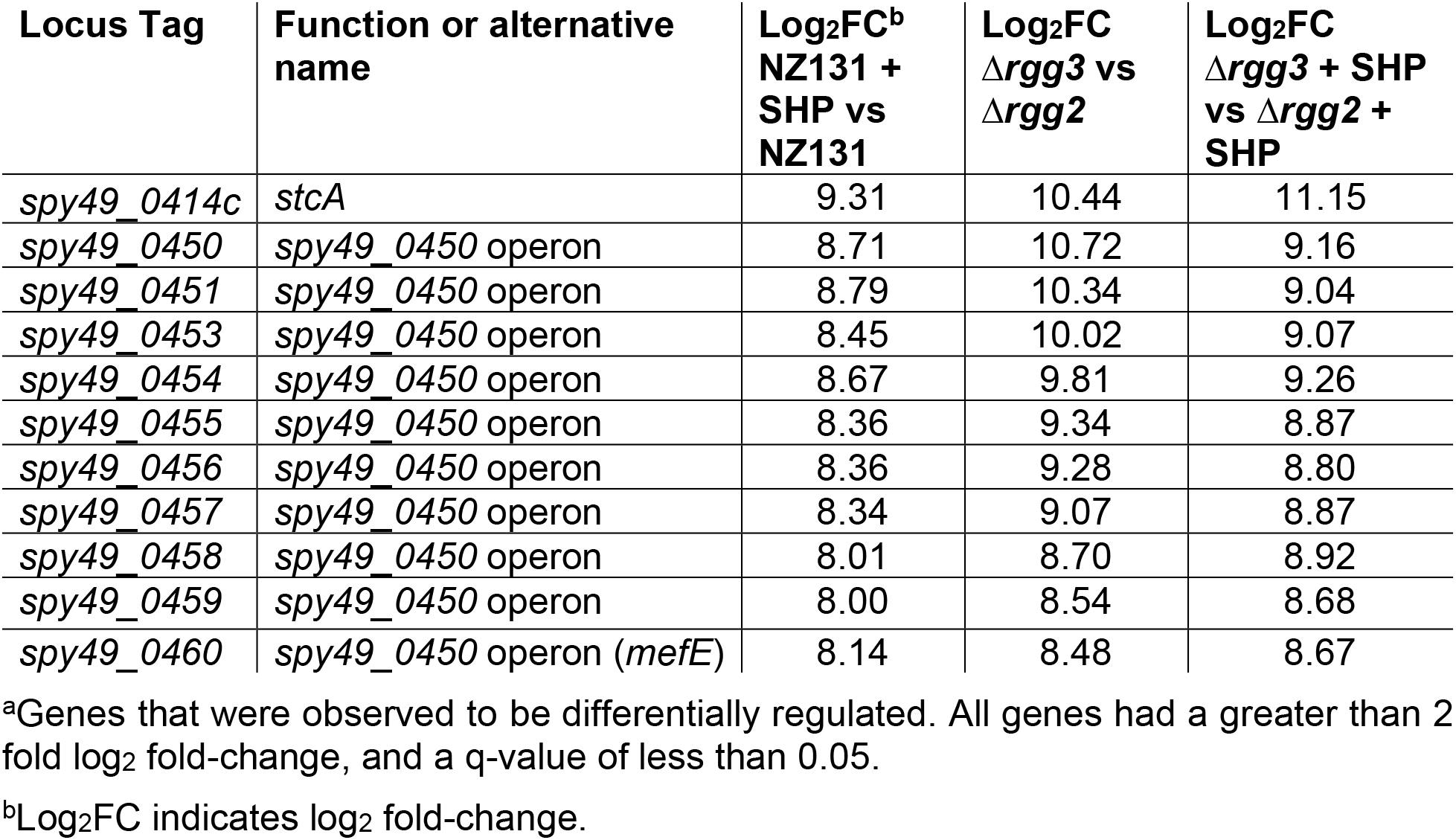
Core regulon of Rgg2/3 QS

| Locus Tag | Function or alternative name | Log<sub>2</sub>FC<sup>b</sup><br>NZ131 +<br>SHP vs<br>NZ131 | Log<sub>2</sub>FC<br>Δ*rgg3* vs<br>Δ*rgg2* | Log<sub>2</sub>FC<br>Δ*rgg3* + SHP<br>vs Δ*rgg2* +<br>SHP |
| --- | --- | --- | --- | --- |
| *spy49_0414c* | *stcA* | 9.31 | 10.44 | 11.15 |
| *spy49_0450* | *spy49_0450* operon | 8.71 | 10.72 | 9.16 |
| *spy49_0451* | *spy49_0450* operon | 8.79 | 10.34 | 9.04 |
| *spy49_0453* | *spy49_0450* operon | 8.45 | 10.02 | 9.07 |
| *spy49_0454* | *spy49_0450* operon | 8.67 | 9.81 | 9.26 |
| *spy49_0455* | *spy49_0450* operon | 8.36 | 9.34 | 8.87 |
| *spy49_0456* | *spy49_0450* operon | 8.36 | 9.28 | 8.80 |
| *spy49_0457* | *spy49_0450* operon | 8.34 | 9.07 | 8.87 |
| *spy49_0458* | *spy49_0450* operon | 8.01 | 8.70 | 8.92 |
| *spy49_0459* | *spy49_0450* operon | 8.00 | 8.54 | 8.68 |
| *spy49_0460* | *spy49_0450* operon (*mefE*) | 8.14 | 8.48 | 8.67 |

<sup>a</sup>Genes that were observed to be differentially regulated. All genes had a greater than 2 fold log<sub>2</sub> fold-change, and a q-value of less than 0.05.
<sup>b</sup>Log<sub>2</sub>FC indicates log<sub>2</sub> fold-change.

It was also of interest to learn whether stimulation of the Δ*rgg2* strain with SHP impacted gene expression, as the hypothesis that Rgg2 is the sole activator of the QS system has not yet been challenged. Comparing mRNA levels of the Δ*rgg2* strain treated with SHP to that of wild type or unstimulated Δ*rgg2* cultures found no significant differences. We applied the same stimulation to the Δ*rgg3* strain. We anticipated only to find an increase in the expression level of the core regulon since the amount of SHP in cultures would be increased. Instead, when we compared stimulated Δ*rgg3* to wildtype unstimulated cultures we found ∼50 additional genes with differential expression patterns (Data Set S1). Some of these genes included *sclB* (*spy49_0830*), *grab* (protein G-related macroglobulin-binding protein, *spy_1080c*), and *pyrR* (*spy49_0650*). A similar number of differentially regulated genes (∼59 additional genes) was observed between SHP-treated Δ*rgg3* and SHP-treated Δ*rgg2* (Fig. 1E-F, Data Set S1). This included genes involved in immune regulation such as *mac* (*spy49_0679*), *emm49* (*spy49_1671c*) and nucleotide metabolism such as *xpt* (*spy49_0887*, xanthine phosphoribosyltransferase), *spy49_0888* (predicted xanthine permease), and *pyrD* (*spy49_1144*).

### SHP-stimulation alters protein production

As phenotypes associated with the Rgg2/3 system include enhanced lysozyme resistance, biofilm formation, and contact-dependent immunosuppression which reflect putative changes to the GAS surface (5, 9, 14), we sought to evaluate proteomic profiles of surface-associated and released proteins from cultures with differentially stimulated Rgg2/3. A published protocol for isolation of proteins from *S. pyogenes* for proteomic evaluation was adapted and performed with tandem mass tag (TMT) labeling in conjunction with LC-MS/MS (22, 23) (Fig. S1A). Cultures of wildtype, Δ*rgg2*, or Δ*rgg3* were grown with or without SHP until late exponential phase (the same point at which transcriptomic evaluations were conducted) when proteins were harvested. Total protein amounts isolated from each fraction did not differ significantly between strains (Fig. S2F). Overall, 404 unique proteins were detected, representing approximately 25% of the theoretical proteome (Fig. S1B). Notably, the number of proteins uniquely detected in cell-free culture supernatants closely matched that reported by Wilk *et. al* (98 proteins total), but fewer proteins were identified in the cell associated fraction (22). Complete lists of detected proteins and statistical analysis are provided in Data Sets S2 and S3.

Members of the core Rgg2/3 regulon (*stcA* and those encoded by genes *0450-0460*) consistently displayed increased amounts in all samples in which quorum sensing was stimulated as compared to unstimulated. A substantially larger number of proteins (114 for cell associated, 66 for secreted fractions, Fig. S1C-D) were altered between SHP-stimulated cultures of Δ*rgg3* and Δ*rgg2*; these changes were not seen between stimulated and unstimulated cultures of wildtype for the cell associated fraction (Fig. 2A). The expanded number of differentially produced proteins in the mutant-strain comparisons are primarily due to the SHP-stimulated Δ*rgg3* condition, as these samples formed a cluster separate from others in the primary dimension when evaluated by principal component analysis (Fig. 3C-D; unstimulated *rgg3* and *rgg2* cultures were not processed due to limited resources). In addition to core-regulon targets being differentially expressed, virulence factors, ribosomal and translation-associated proteins, and nucleotide biosynthesis protein levels were altered in the cell-associated fraction (Fig. 2B, Data Set S2). A similar trend in ribosomal and translation-related proteins was observed when comparing the stimulated Δ*rgg3* condition to wild-type with or without SHP (Fig. S2A-B), which was not observed in stimulated Δ*rgg2* versus wild-type (Fig. S2C-D).

Since previous work has found that up to 60% of proteins may display poor correlation between cellular concentrations and their corresponding RNA transcripts (24, 25), we attempted to validate RNA-seq and proteomic results from additional biological replicates using targeted RT-PCR and western blotting. Transcripts of the primary Rgg2/3 targets *spy49_0450* and *stcA* were confirmed as substantially upregulated (Fig. 4A-B). Several additional subjects were selected based on their roles as virulence factors or differential expression pattern in both the RNA-seq and proteomic datasets. These included: *slo, upp, spy49_0877, plr, prtF, ska, sfbX49*, and *sclB* (Fig. 4, Fig. S3). Half of these targets (*slo, upp, prtF, sfbX49*) displayed transcriptional changes between conditions, but the others could not be statistically validated (*spy49_0877, plr, ska, sclB*) (Fig. 4, S3).

**FIGURE 4:**
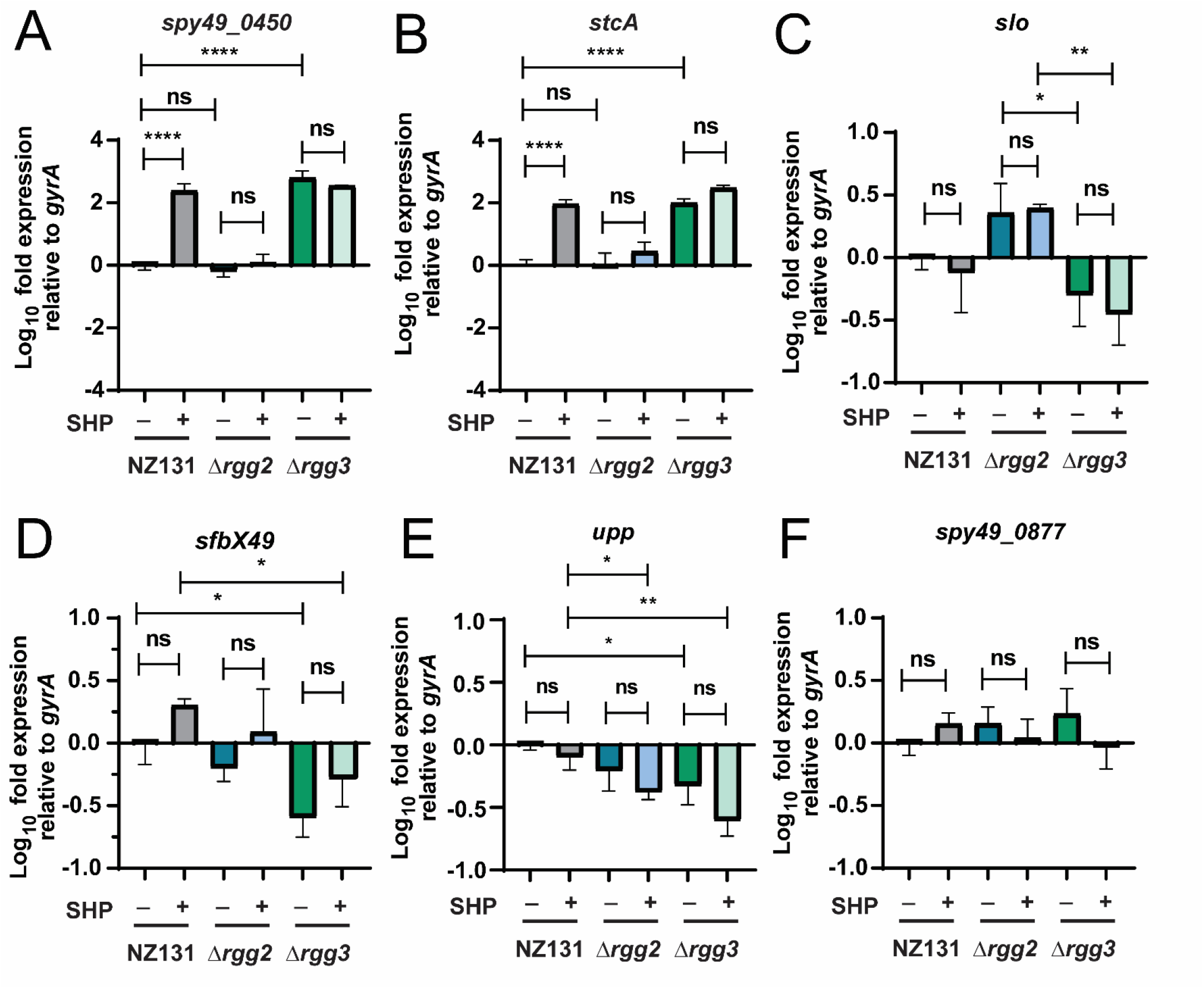
qRT-PCR results verifying RNA-seq and examining if proteomics targets have altered transcript levels. Wild-type NZ131, Δ*rgg2*, and Δ*rgg3* strains were grown in biological triplicate with or without 100 nM SHP peptide (indicated by + or – sign below each graph) and RNA was harvested late exponential phase. RNA was processed for qRT-PCR and transcript levels were determine relative to the *gyrA* reference gene. Significance of transcript level changes were determined using a One-way ANOVA with Multiple Comparisons Post-test. *, P < 0.05; **, P < 0.005; ****, P < 0.0005; ns, non-significant. For further experimental details, see *Materials and Methods*. A) Relative transcript levels of *spy49_0450 (aroE.2)*. B) Relative transcript levels of *stcA* (*spy49_0414c*). C) Relative transcript levels of *slo* (*spy49_0146*). D) Relative transcript levels of *sfbX49* (*spy49_1683c*). E) Relative transcript levels of *upp* (*spy49_0322*). F) Relative transcript levels of *spy49_0877*.

To evaluate changes in protein levels, we constructed sfGFP fusions to two proteins of interest: AroE.2 (Spy49_0450) and Spy49_0877. AroE.2 was only detectable in the stimulated wildtype or in Δ*rgg3* but not in unstimulated wildtype or either Δ*rgg2* condition (Fig. 5A, 5C). For Spy49_0877, the SHP-induced change observed in the proteomics dataset was slight but significant (∼1.3 ratio increase in expression; Data Set S2); however, the fusion to sfGFP showed no significant differences in levels between strains or conditions (Fig. 5B, 5D). Taken together, validation was confirmed for the core targets of Rgg2/3 regulation, whereas small changes observed in transcriptomic and proteomic datasets are likely stochastic fluctuations resulting from apparent stress induced responses.

**FIGURE 5:**
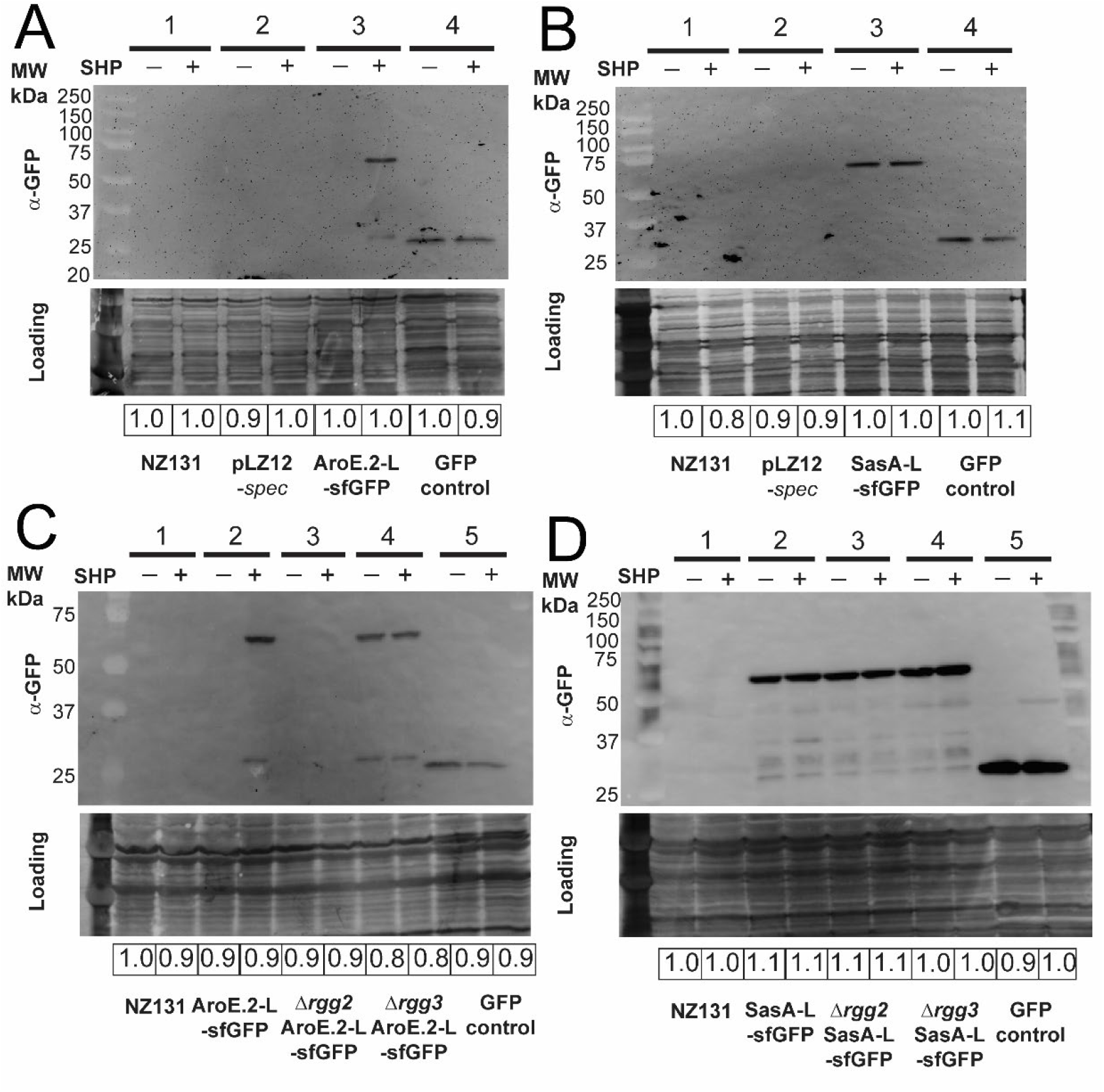
Western blots of select proteins validating proteomics data. Blots were probed with α-GFP. India Ink stain was used to assess gel loading and relative loading is shown below for each blot. GFP expressed from a plasmid in GAS validated antibody specificity. Shown beside blots are the corresponding molecular weights of the ladder. +/- indicates cultures with or without 100 nM SHP. For further details, see *Materials and Methods*. All experiments were performed three times. A) Western blot examining expression of AroE.2-L-sfGFP in NZ131 wild-type background, +/- SHP peptide. 1: NZ131 wild-type; 2: NZ131 pLZ12-*spec*; 3: NZ131 pBER40 (pLZ12-*spec*-*aroE.2*-*L*-*sfGFP shp3*(ATG→GGG)); 4: GFP control (HSC5 pFED630). Expected protein sizes in kDa: AroE.2-L-sfGFP 63.5; sfGFP 27.0; GFP 26.8. B) Western blot examining expression of SasA-L-sfGFP in NZ131 wild-type background, +/- SHP peptide. 1: NZ131 wild-type; 2: NZ131 pLZ12-*spec*; 3: NZ131 pBER41 (pLZ12-*spec*-*sasA*-*L*-*sfGFP*); 4: GFP control (HSC5 pFED630). Expected protein sizes in kDa: SasA-L-sfGFP 54.3, sfGFP 27.0, GFP 26.8. C) Western blot examining expression of AroE.2-L-sfGFP in NZ131 wild-type, Δ*rgg2*, or Δ*rgg3* backgrounds +/- SHP peptide. 1: NZ131 wild-type; 2: NZ131 pBER40 (pLZ12-*spec*-*aroE.2*-*L*-*sfGFP shp3*(ATG→GGG)); 3: NZ131 Δ*rgg2* pBER40; 4: NZ131 Δ*rgg3* pBER40; 5: GFP control (HSC5 pFED630). Expected protein sizes in kDa: AroE.2-L-sfGFP 63.5, sfGFP 27.0, GFP 26.8. D) Western blot examining expression of SasA-L-sfGFP in NZ131 wild-type, Δ*rgg2*, or Δ*rgg3* backgrounds, +/- SHP peptide. 1: NZ131 wild-type; 2: NZ131 pBER41 (pLZ12-*spec*-*sasA*-*L*-*sfGFP*); 3: NZ131 Δ*rgg2* pBER41; 4: NZ131 Δ*rgg3* pBER41; 5: GFP control (HSC5 pFED630). Expected protein sizes in kDa: SasA-L-sfGFP 54.3, sfGFP 27.0, GFP 26.8.

### Rgg2/3 stimulation impacts growth rate and has a link to stringent response

Several proteins displaying altered levels in the proteomics dataset are involved in nucleotide biosynthesis, most notably those involved in stringent response or purine biosynthesis (PurA, PurC, PurL, PurK, PurH, GuaB, Spy49_0877) (Data Set S2). As regulatory links have been documented between some quorum sensing systems and the stringent response (26, 27), we sought to test a possible connection between the stress-response system and Rgg2/3. In prior studies, Steiner and Malke found that RelA (*spy49_1632c*) was responsible for the majority of (p)ppGpp accumulation under amino acid starvation in *S. pyogenes*, but they also identified two additional putative alarmone synthases (*spy49_0877* and *spy49_0687* in NZ131, *spy_1125* and *spy_0873* in SF370, respectively) (28, 29). RelA is predicted to have two enzymatic activities, serving as a bifunctional synthetase and hydrolase, whereas Spy49_0877 and Spy49_0687 are predicted to encode small alarmone synthases (SASs) that we have named *sasA* and *sasB*, respectively, based on orthologous genes in *B. subtilis* (30, 31). As *sasA* and *sasB* have remained untested in *S. pyogenes*, we took the opportunity to construct deletions of each synthase gene, as well as double and triple knockout combinations with *relA* (Fig. 6). Growth-rate phenotypes were evaluated for each mutant and were compared with wildtype, *Δrgg2*, and *Δrgg3* strains upon stimulation with SHP. Doubling times of SHP-stimulated NZ131 were shorter than unstimulated (36 ± 1 min unstimulated vs. 45 ± 2 min stimulated) and addition of a reversed-sequence SHP peptide did not delay growth (Table S2). The Δ*rgg2* strain was unresponsive to SHP, growing at the same rate as unstimulated wildtype, whereas SHP-supplemented Δ*rgg3* grew substantially slower when treated with SHP (39 ± 4 min unstimulated vs. 57 ± 2 min stimulated), consistent with the idea that excess SHP poses a stress to cells (Table S2, Fig. 6B). In observing growth rates of the guanosine-phosphate synthases, SHP stimulation had little effect on the *sasA* and *sasB* deletions but significantly slowed the growth of the *relA* deletion (Fig. 6C, Table S2). Complementing *relA* restored its growth to a wildtype rate (Fig. S4A). We also observed that presence of a second copy of *sasA* or *relA* but not *sasB* abrogated the SHP-dependent increase in doubling time compared to wild-type, although a slight increase in doubling time was still observed for these strains but was non-significant (Table S2, Fig. S4A-C). We then examined growth of double and triple knockout strains. Although growth rates were unaffected for double-knockout strains (Δ*relA* Δ*sasA,* Δ*relA* Δ*sasB,* and Δ*sasA* Δ*sasB*) (Table S2, Fig. 6D-F), the triple-knockout (Δ*relA* Δ*sasA* Δ*sasB*) displayed a longer doubling time for both stimulated and unstimulated conditions (62 ± 8 min unstimulated and 72 ± 5 min stimulated) (Table S2, Fig. 6D).

**FIGURE 6:**
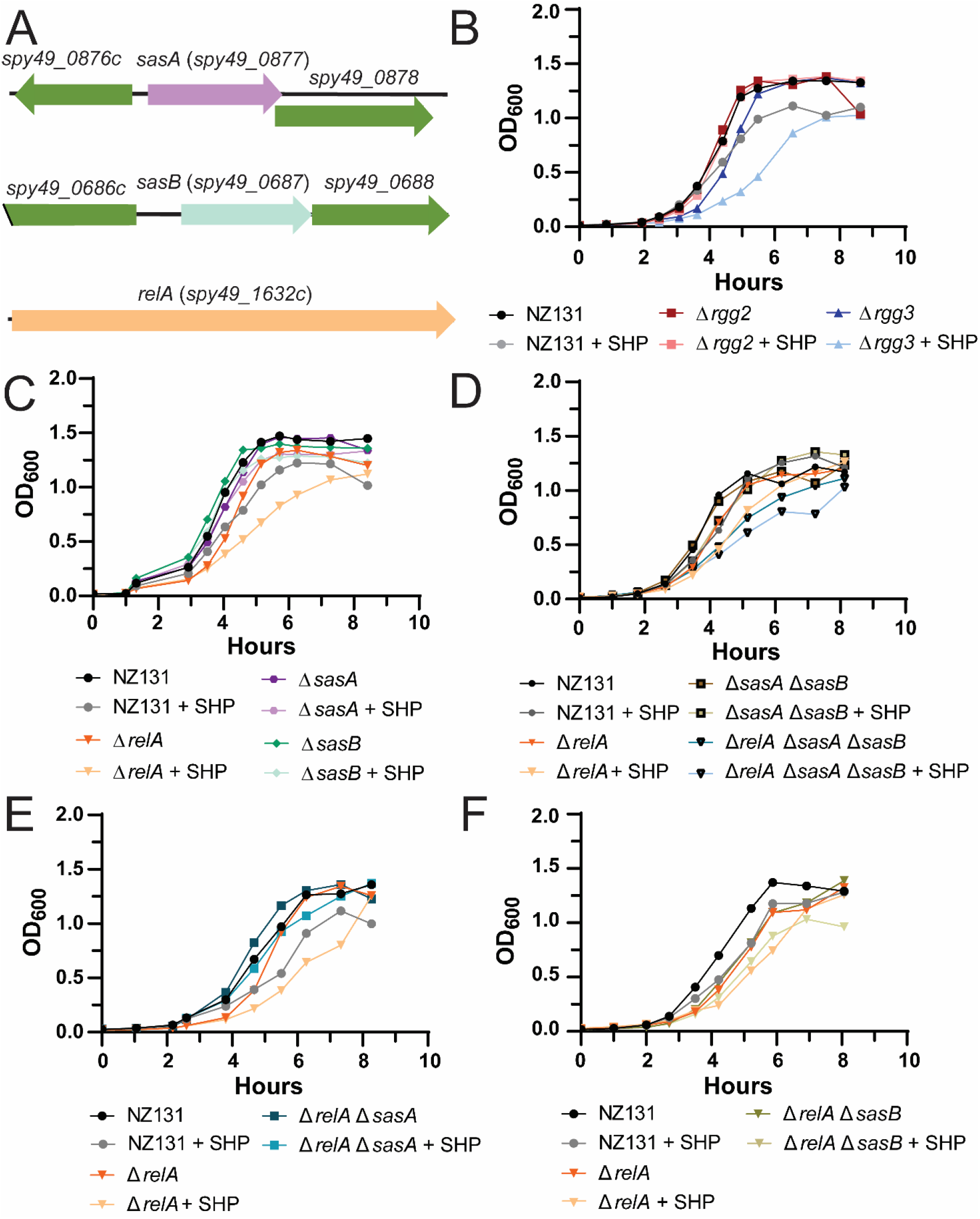
Examination of the connection between the Rgg2/3 system and stringent response. All experiments were performed a minimum of three times. A) Genetic loci of *sasA* (*spy49_0877*), *sasB* (*spy49_0687*), and *relA* (*spy49_1632c*), shown to relative scale. B) Growth curve of wild-type NZ131, Δ*rgg2*, and Δ*rgg3* with or without 100 nM SHP peptide. Strains and conditions are indicated in the legend below the graph. C) Growth curve of wild-type NZ131, *ΔrelA*, Δ*sasA*, and Δ*sasB* with or without 100 nM SHP peptide. Strains and conditions are indicated in the legend below the graph. D) Growth curve of single deletion of *relA*, compared to double and triple deletions. Wild-type NZ131, Δ*relA*, Δ*sasA* Δ*sasB*, and Δ*relA* Δ*sasA* Δ*sasB* were grown with or without 100 nM SHP peptide. Strains and conditions are indicated in the legend below the graph. E) Growth curve of single deletion of *relA* compared to double deletion strain. NZ131, Δ*relA*, and Δ*relA* Δ*sasA* were grown with or without 100 nM SHP peptide. Strains and conditions are indicated in the legend below the graph. F) Growth curve of single deletion of *relA* compared to double *relA sasB* deletion. NZ131, Δ*relA*, and Δ*relA* Δ*sasB* were grown with or without 100 nM SHP peptide. Strains and conditions are indicated in the legend below the graph.

Given the compounding effects on growth rate seen in mutants stimulated with SHP, we investigated whether the two regulatory systems (QS and stringent response) had reciprocal regulatory impacts on one another. While transcriptional expression of *sasA* and *sasB* were unaffected by Rgg2/3 activity, as determined by luciferase reporters (Fig. S4E-F, unfortunately we were unable to generate a P*_relA_*-*luxAB* reporter), the impact of alarmone synthase mutants on QS activity was apparent under certain conditions. Previous studies (14, 32) have demonstrated that induction of Rgg2/3 is autonomous in the presence of mannose, as can be observed using luciferase fusions to the *shp* promoter (Fig 7). Mannose-dependent induction was eliminated in the *relA* mutant and partially incapacitated in the *sasA* mutant, but not affected by *sasB*.

**FIGURE 7:**
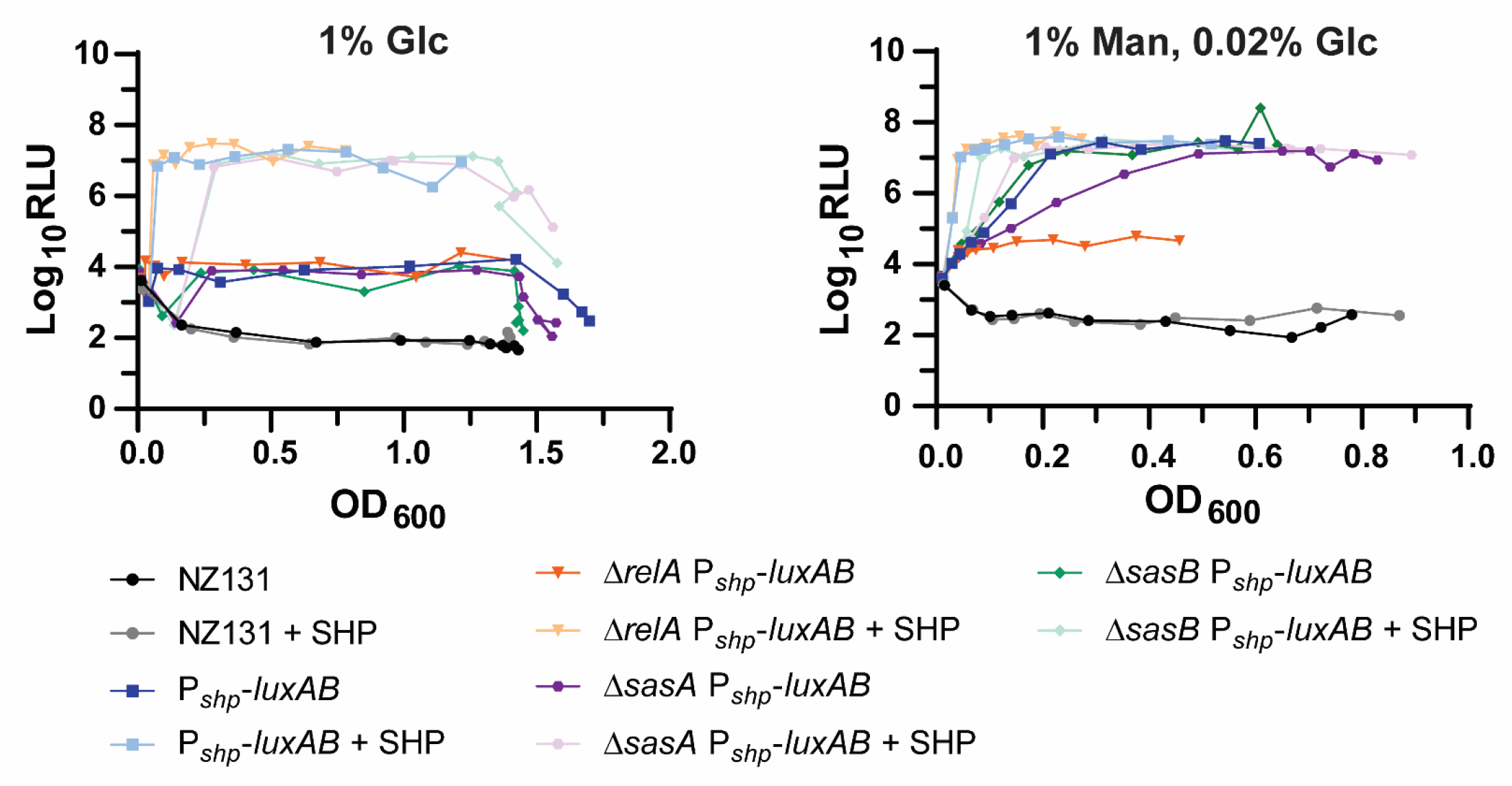
Effect of deletion of Rel homologs on P*shp* activity in different carbon sources. Luciferase assays of P*shp* reporter strains in the presence of 1% glucose (1% Glc) or 1% mannose supplemented with 0.02% glucose (1% Man, 0.02% Glc). The assay in 1% Glc was performed twice with similar results, whereas the assay in 1% Man, 0.02% Glc was performed three times. Wild-type, P*shp*-*luxAB*, Δ*relA* P*shp*-*luxAB,* Δ*sasA* P*shp*-*luxAB*, and Δ*sasB* P*shp*-*luxAB* were grown in the above conditions with or without 100 nM SHP peptide. Luciferase activity was monitored as outlined in *Materials and Methods*. Strains and conditions are indicated in the legend below the graph.

Finally, to observe relative GTP and (p)ppGpp levels upon induction or disruption of the Rgg2/3 system, thin-layer-chromatography (TLC) was conducted following incorporation of radiolabeled ^32^P-orthophosphate during culture growth. Consistent with the modest increases in nucleoside synthases seen from the proteomic dataset, small increases in levels of ppGpp and GTP were observed in wildtype induced with SHP and in the Δ*rgg3* strain in comparison to Δ*rgg2* (Fig. 8). Given that nucleotide levels were altered and that Δ*relA* was unable to respond to mannose conditions, we conclude that an indirect connection between the Rgg2/3 system and stringent response exists, possibly as an adaptation to exposure of high concentrations of the hydrophobic SHP peptide, to high requirements of branched-chain amino acids during SHP production, or to dysregulation of nutritional status monitoring.

**FIGURE 8:**
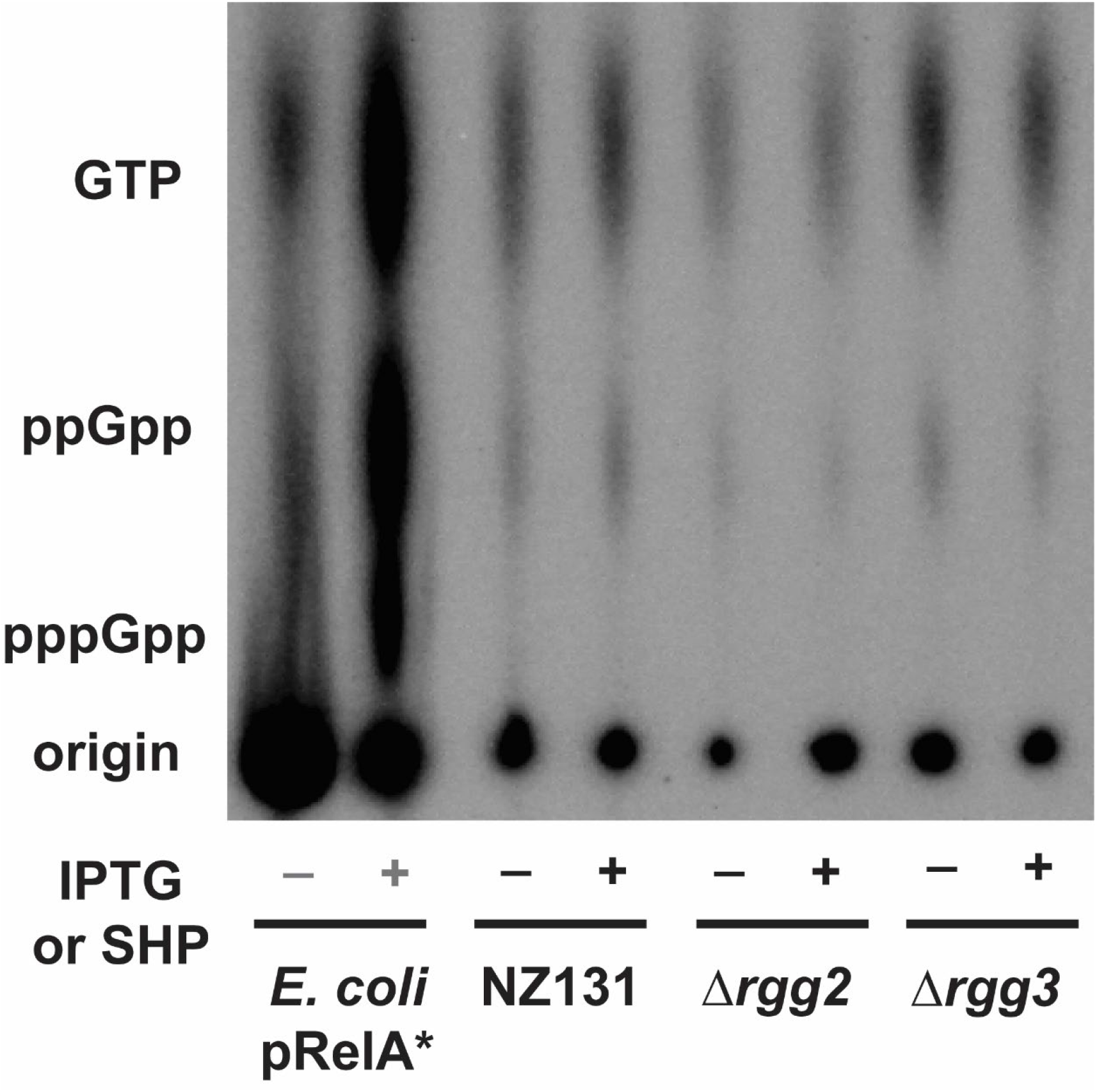
Impact of SHP induction and disruption of the Rgg2/3 system on nucleotide production. Thin layer chromatography (TLC) examining nucleotide levels in cells grown in MOPS-CDM with 150 µCi/mL ^32^P-orthophosphate. As a control, an IPTG inducible constitutive form of RelA from *E. coli* was run to provide standards for GTP, ppGpp, and pppGpp production, - indicates no IPTG added, + indicates 100 mM IPTG was added. Wild-type NZ131 and derivative strains were grown with or without 100 nM SHP peptide, as indicated by the -/+ below the TLC. Labels beside the TLC indicate the location of GTP, ppGpp, and pppGpp as based on the IPTG induced *E. coli* pRelA* standard. This experiment was performed three times. Image shown was evenly adjusted for contrast. For further experimental details, see *Materials and Methods*.

**FIGURE 9:**
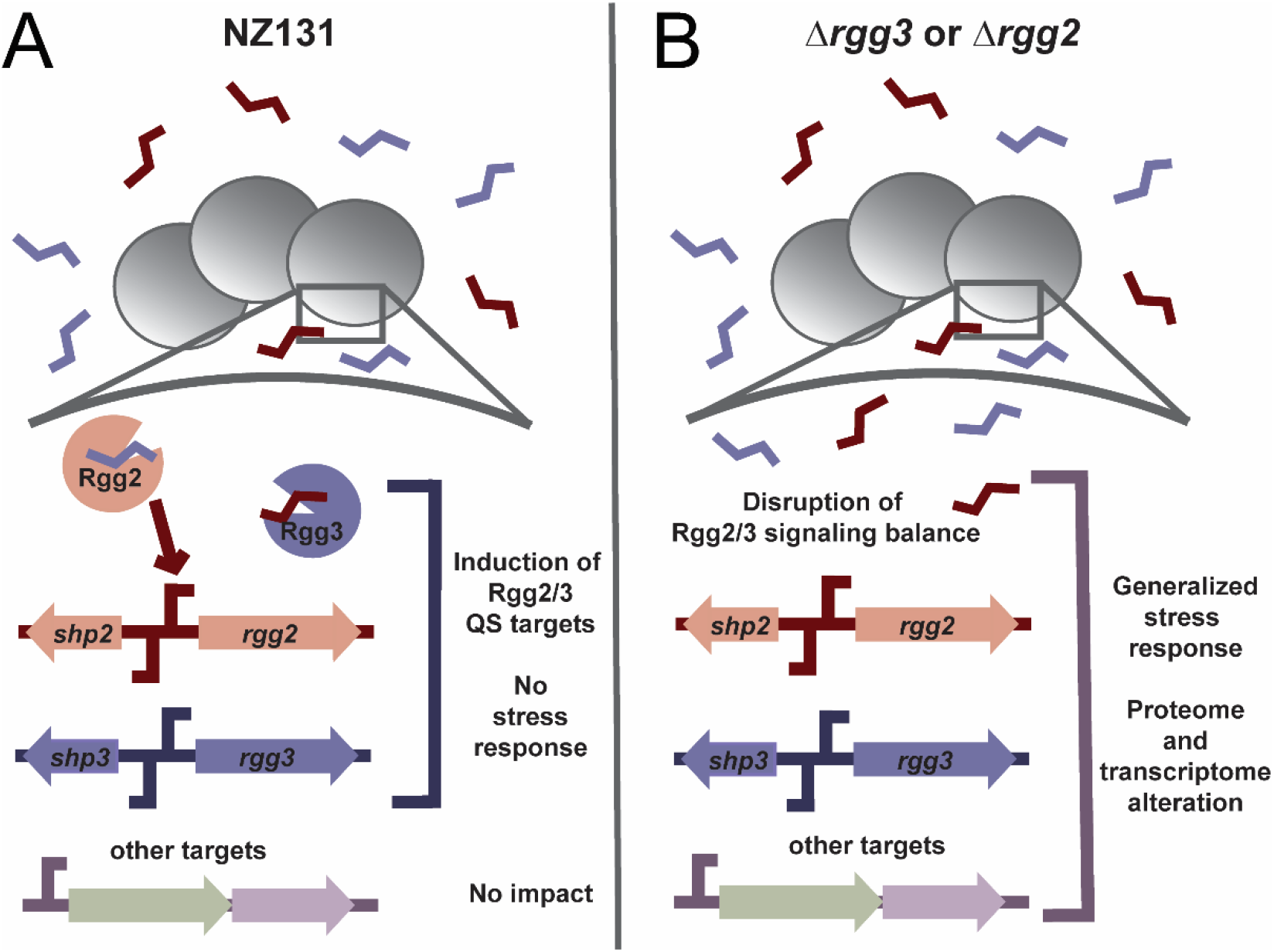
Model illustrating the impact of Rgg2/3 deletion. Panel A illustrates what occurs when SHPs are added in a wild-type NZ131 background. Rgg3 binds SHPs and is released from target promoters, while Rgg2 binds SHPs and activates target promoters. This includes the SHP2/Rgg2 and SHP3/Rgg3 loci, as well as others such as the *spy49_0450* operon and *stcA*. Normal Rgg2/3 QS occurs and a small number of target transcripts are affected. Panel B illustrates what occurs in the absence of Rgg2 or Rgg3, primarily Rgg3. Disruption of the system leads to an imbalance in QS activation. This results in a generalized stress response that alters the transcriptome and proteome of NZ131 cells.

## DISCUSSION

The objective of this study was to determine the genome-wide impact on gene and protein expression as a function of signaling by the Rgg2/3 quorum sensing system in *S. pyogenes*. Our ongoing mechanistic understanding of gene regulation governed by two transcriptional regulators, Rgg2 and Rgg3, was further validated through analysis of responses to SHP pheromone and in use of null mutants. Stimulation of wildtype with SHP confirmed that two operons, whose transcription start sites include the Rgg2/3 bindings sequence, are the primary and direct targets of this regulatory network. This is not surprising, as previous work identified sequences required for Rgg2/3 binding are proximal to the *shp2* and *shp3* promoters and needed for robust regulation by the two transcriptional regulators in response to pheromone (17). The sequences occur only twice in the *S. pyogenes* genome, at the primary targets of the system, and are found conserved in other streptococcal species, including *S*. *agalactiae*, *S*. *dysgalactiae*, and *S*. *pneumoniae* where orthologs of Rgg2 or Rgg3 are present (33). As deletion of *rgg3* removes the repressive impact on target promoters, and therefore increases production of SHP pheromones, Δ*rgg3* mutants express Rgg2/3 target genes in a manner highly similar to that of wild-type cultures of NZ131 responding to exogenous SHP. Conversely, mutants of *rgg2* no longer have the transcriptional activator necessary for promoter induction; therefore, Δ*rgg2* mutants are incapable of turning the circuit on, even with exogenously supplied SHP, as the results presented herein substantiate. The equivocation of Δ*rgg3* as being QS-ON, and Δ*rgg2* as QS-OFF is upheld by results presented here.

Recently we employed Δ*rgg2* and Δ*rgg3* mutants generated in three different GAS serotypes and studied their ability to colonize the murine upper respiratory tract. Though the three serotypes colonized to different extents and for different lengths of time, in each case the Δ*rgg3* mutant (QS-ON) colonized as well or to higher numbers than wildtype. The Δ*rgg2* mutant (QS-OFF) were all poor colonizers, indicating the Rgg2/3 system contributes colonization ability (21). Likewise, QS-ON and QS-OFF mutants held opposite abilities at inhibiting immune activation of macrophages. The QS-ON cells (Δ*rgg3*, or wildtype stimulated with SHP) suppressed macrophage activation as reported by NF-kB activity (9). Thus, the genes encoded in the core regulon should be the focus of future studies aimed at determining the mechanisms providing for tissue colonization and immunosuppression.

However, our analyses also revealed that cultures of Δ*rgg3* (which produce SHP endogenously to levels at least matching the functional equivalent of exogenously supplied 100 nM to wildtype; compare fold changes of the *0450* operon genes between Fig. 1A and C), when stimulated further by exogenously supplied 100 nM SHP, had a more severe impact on gene and protein expression, beyond changes to the core regulon. The increased differential in expression and in numbers of genes and proteins, when compared to WT, WT+SHP or Δ*rgg2*+SHP, was primarily due to changes seen in the Δ*rgg3*+SHP samples, best visualized by principle component analysis (Fig. 1E-F, Fig. 2B-D, Fig. 3, Fig. S1). This extreme condition affected the expression of important virulence factors (*prtF*, *sfbX49*, *slo*) and proteins involved in translation and nucleotide biosynthesis, and we suspected this impact was due to excessive levels of the hydrophobic SHP peptide. The observed increase in SasA protein amounts and a general downshift in translation-related proteins in SHP stimulated Δ*rgg3* cultures led us to hypothesize an interplay between the Rgg2/3 system and stringent response. Stringent response induction in *E. coli* and *B. subtilis* has been associated with a large transcriptional shift (34, 35), which we found to occur in SHP-induced Δ*rgg3*. To test this hypothesis, we constructed individual and combinatorial gene deletions of the three enzymes predicted or demonstrated as involved in stringent response (28). While minimal effects on Rgg2/3 signaling were seen in knockouts of *sasA* and *sasB,* the Δ*relA* strain enhanced the SHP-induced growth phenotype, and more remarkably, blocked the ability of mannose to induce Rgg2/3 signaling. Our lab recently demonstrated that Rgg2/3 QS induction in mannose is connected to the phosphorylation status of HPr, specifically via its impact on Mga phosphorylation (32). In *E. coli,* the activity of the primary ppGpp hydrolase, SpoT, is impacted by HPr phosphorylation status and carbon source via its interaction with the protein Rsd (36). Although there is no homolog of *rsd* in the *S. pyogenes* genome, a similar connection between HPr phosphorylation status and RelA hydrolytic activity could be envisioned and would provide an indirect connection between mannose induction of Rgg2/3 QS and stringent response.

We also examined if any change in GTP or (p)ppGpp levels occurred upon stimulation or disruption of the Rgg2/3 system. Curiously, SHP pheromones are rich in branched chain amino acids (SHP2: MKKVNKALLFTLIMDILIIVGG; SHP3: MKKISKFLPILILAMDIIIIVGG) and we have suspected that excessive induction of the system (as would be the case in Δ*rgg3*+SHP conditions) would lead to exaggerated expression and translation of SHPs, possibly depleting branched-chain amino acid (BCAA) pools, and potentially converting RelA from hydrolase to synthase activity via the interaction of GTP with CodY (37–39). Alternatively, if CodY is unresponsive to GTP levels such as observed in *S. pneumoniae* and *S. mutans*, perhaps nucleotide levels are affected by inhibition of (p)ppGpp hydrolysis via BCAA depletion (40, 41). This phenotype has been observed for Rel from *Rhodobacter capsulatus* and BCAAs bind Rel proteins in some Gram-positive species (42). This has not been determined for *S. pyogenes*, but examination of *codY* and *relA* regulons in *S. pyogenes* indicated little overlap, hinting that the secondary hypothesis might be more likely (43).

Though only a slight difference in basal GTP and ppGpp levels was observed, and no overt induction of pppGpp was found under these conditions, these emerging signs of stress and delayed growth are consistent with the idea that excessive induction of the Rgg2/3 system has a negative impact on cellular physiology. We conclude that whatever feedback is occurring between the Rgg2/3 system and stringent response is likely indirect and seen under artificial conditions. Nevertheless, the observed impact we document should be considered in future laboratory studies in which signaling systems are subject to exogenous and genetic manipulations. In total, this work provides a clearer picture of the impact of the Rgg2/3 system on the proteome and transcriptome of *S. pyogenes* and provides a framework for further understanding how this QS system affects cell physiology.

## MATERIALS AND METHODS

### Bacterial strains, plasmids, and growth conditions

Bacterial strains and plasmids used in this study are listed in Table S1. *S. pyogenes* strains were derived from NZ131 or HSC5 (16, 44) and were grown on Todd-Hewitt plates (TH, BD Biosciences) with 1.4 Bacto-Agar (BD Biosciences) and 0.2% yeast (BD Bioscienes), in Todd-Hewitt broth with 0.2% yeast (THY), in chemically-defined medium (CDM) plus 1% glucose or 1% mannose with 0.02% glucose, or in MOPS-CDM (see next section) at 37°C. The components and recipe for CDM used were as described previously (14, 16). *E. coli* strains were grown on Luria-Bertani (BD Biosciences) plates with 1.4% Bacto-Agar at 37°C or in Luria-Bertani (LB) broth with shaking at 175 rpm. When required, ampicillin (100 µg/mL, *E. coli* only), spectinomycin (100-200 µg/mL), erythromycin (0.5 µg/mL for *S. pyogenes*, 500 µg/mL for *E. coli*), kanamycin (50-100 µg/mL), chloramphenicol (3-6 µg/mL) were added to *S. pyogenes* or *E. coli* culture media.

### Preparation of MOPS-CDM

MOPS-CDM recipe was based on Kazmierczak *et. al*, 2009 (45). In brief: MOPS-CDM replaces potassium phosphate and sodium phosphate in the original CDM recipe with 40 mM MOPS, 4 mM Tricine, 0.28 mM K_2_SO_4_, and 50 mM NaCl. All other components were kept the same as in the original CDM recipe with 1% glucose added (16).

### Construction of plasmids in *E. coli*

To construct plasmids, one of the four following methods was used: restriction digest and ligation, Gibson assembly, a combination of restriction digest and Gibson assembly, or quick-change mutagenesis. Method used for construction of each plasmid, primers and additional information is listed in Table S1. All plasmids were propagated in DH5α cells and extracted using a GeneJET Plasmid MiniPrep Kit (Thermo-Fisher) according to the manufacturer’s instructions. All PCR amplifications were carried out with Phusion High-Fidelity DNA Polymerase (New England Biolabs, NEB), and resulting fragments were purified with DNA Clean & Concentrator kit (Zymo Research). Restriction digests (NEB) were carried out according to the manufacturer’s instructions. All final plasmid constructs were electroporated into *E. coli* DH5α using a Bio-Rad Gene Pulser II Electroporation system (Bio-Rad) with the following parameters: 2.0 V, 200 Ω, 250 µF. Constructed plasmids were confirmed by restriction digest, PCR and/or Sanger sequencing.

To construct plasmids by restriction digest and ligation, PCR obtained linear fragments or plasmid vectors were digested with restriction enzymes as indicated in Table S1. Resulting insert and vectors were purified and ligated together using T4 DNA ligase (NEB) according to the manufacturer’s instructions. To construct plasmids by Gibson assembly, plasmid backbone and inserts were obtained from templates via PCR. Resulting fragments were purified and assembled via Gibson assembly using NEBuilder HiFi DNA Assembly MasterMix (NEB). To construct plasmids by combination of restriction digest and Gibson assembly, inserts for plasmids were obtained via PCR and assembled via Gibson assembly. Resulting inserts and plasmid vectors were digested with restriction enzymes as outlined in Table S1 and ligated together with T4 DNA ligase. For pBER18, an intermediate plasmid was required for construction. The 3’ flanking region near *spy49_0877* was amplified, digested, and ligated into pFED760. The resulting intermediate plasmid and the 5’ flanking region near *spy49_0877* was then PCR amplified and assembled via Gibson assembly to obtain the final pBER18 plasmid. For quick-change mutagenesis, required quick change primers as indicated in Table S1 were used, following the protocol for Phusion Site-Directed Mutagenesis from Thermo-Fisher Scientific. The resulting linearized quick-change plasmid was then digested with DpnI (NEB) to eliminate the parent plasmid and ligated using T4 DNA ligase.

### Construction of *S. pyogenes* strains with markerless deletions

To construct markerless deletions of *relA*, *sasA*, *sasB*, and combinations thereof, the following procedure based on the protocol for Gene Replacement by Allelic Exchange published by Le Breton and McIver was used (46). All strains and their derivatives are listed in Table S1. Required knockout plasmids (pBER16, pBER17, or pBER18) were propagated in *E. coli* DH5α and extracted using a GeneJET Plasmid MiniPrep Kit. 500 ng of purified plasmids were then electroporated into NZ131 using the following parameters: 1.75 V, 400 Ω, 250 µF. After electroporation, cells were allowed to recover in THY at 30°C for 2 hours, and were then plated on THY + Erm plates and incubated for two days at 30°C. Resulting colonies were patched on THY + Erm plates to confirm resistance and growth at 30°C and to confirm inability to grow at 37°C. Colonies were then grown overnight at 30°C with Erm, and additionally stored as ‘unintegrated plasmid’ stocks with 20% glycerol at −70°C. Resulting stocks were diluted 1:100 into fresh THY + Erm and grown at 30°C for 2 hours. After 2 hours, cultures were shifted to 37°C and grown for 3 hours to allow the plasmid to integrate into the chromosome. Cultures were consecutively diluted 1:10 into THY, plated on THY + Erm, and incubated at 37°C overnight. Individual colonies were streaked out THY + Erm plates and incubated at 37°C again. Resulting colonies were PCR confirmed for plasmid integration, and overnight cultures from these colonies were diluted 1:100 into fresh THY (without Erm). Cultures were passaged 1:100 into fresh THY 5 times, consecutively diluted 1:10 and plated on THY plates. Individual colonies were patched on THY and THY + Erm plates and confirmed for loss of Erm^R^ at 37°C. Resulting Erm^S^ colonies were then grown overnight at 37°C in THY and stored as final markerless knockout stocks at −70°C with 20% glycerol. Genomic DNA from resulting strains was then isolated and strains were confirmed by PCR and/or sequencing.

### Construction of other *S. pyogenes* strains

To construct other *S. pyogenes* strains in NZ131 or HSC5, 500 ng of required plasmid (Table S1) was electroporated into the parent strain as previously mentioned. Resulting strains were single-colony isolated, stored as 20% glycerol stocks at −70°C and confirmed by PCR and/or sequencing. All strains and their derivatives are listed in Table S1.

### Synthesis of SHP3-C8 (SHP) and reverse SHP3-C8 (revSHP) peptides

Synthetic peptides were purchased from ABClonal (Woburn, MA) and have been described previously (16, 17). Purities and preparations used in assays were greater than 70%. All peptides were reconstituted as 10 mM stocks in DMSO and stored in aliquots at −20°C. Subsequent dilutions for working stocks (100 µM) were made in DMSO and stored at − 20°C.

### Isolation of RNA from bacterial cultures for qRT-PCR and RNA-seq

Glycerol stocks of bacterial strains at −70°C were inoculated into THY broth and incubated overnight at 37°C. The next morning, strains were diluted 1:100 into 6 mL of prewarmed CDM and grown at 37°C, with OD_600_ observed every 45 min to 1 hour with a GENESYS 30 Vis spectrophotometer (Thermo-Fisher). When strains reached OD_600_ ∼0.05-0.1, 100 nM SHP peptide was added to required tubes. Cultures were grown until OD_600_ reached ∼0.8- 1.0. At this time, 1.5 mL of cultures were harvested and centrifuged at 2700×*g* for 10 min. Supernatant was discarded and cells were resuspended in 750 µL RNA-later solution (Thermo-Fisher, AM7020), and allowed to sit for 10 min at room temperature. Cells were then centrifuged at 21,130×*g* for 1 min, supernatant was discarded, and cells were frozen at −70°C. RNA was extracted from frozen cell pellets using the RiboPure-Bacteria kit (Thermo-Fisher, AM1925) according to manufacturer’s instructions. Samples were DNAse treated with DNAse I from the RiboPure-Bacteria kit according to the manufacturer’s instructions, and RNA concentration and quality was assessed via spectrophotometer (Nanodrop 1000 Spectrophotometer). Samples were additionally run on a 0.9% agarose gel via electrophoresis to confirm presence of 23S and 16S rRNA bands, to confirm samples had not degraded.

### RNA quality determination, preparation of cDNA libraries for DNA sequencing, and total RNA-seq

RNA-seq was conducted in the Northwestern University NUSeq Core Facility. Briefly, RNA quantity was determined via Qubit fluorometer. Total RNA examples were also checked for fragment sizing using a Bioanalyzer 2100 (Agilent). The Illumina Stranded Total RNA Library Preparation Kit (Illumina) was used to prepare sequencing libraries from 500 ng of total RNA samples, according to manufacturer’s instructions. This procedure includes rRNA depletion with RiboZero Gold, cDNA synthesis, 3’ end adenylation, adapter ligation, library PCR amplification and validation. lllumina HiSeq 4000 Sequencer (Illumina) was used to sequence the libraries with the production of single-end, 50 bp reads.

### RNA-seq analysis

The raw RNA sequencing data was processed using the RStudio platform (RStudio 1.4, Boston, MA), with the “Bioconductor” (47, 48) packages “Rsubread” (49) and “Rsamtools”(50) to align RNA-sequencing reads to the *S. pyogenes* strain NZ131 reference genome (GenBank assembly accession: GCA_000018125.1). Annotated feature counts were then produced from the aligned read data, which was subsequently filtered and normalized to generate the final count data for differential expression (DE) analysis using the “edgeR” (51, 52) and “qvalue” (53) packages from “Bioconductor”. Venn diagrams illustrating sample analysis were created using the BioVenn online platform (54).

### Preparation of cDNA for qRT-PCR and qRT-PCR experiments

Total RNA from *S. pyogenes* strains was used to prepare cDNA using the Superscript III First-Strand Synthesis System (Thermo-Fisher) according to the manufacturer’s instructions, including treatment with RNase H. Random hexamer primers were used for cDNA synthesis. All qRT-PCR primers used are listed in Table S1. cDNA was diluted 1:10 and qRT-PCR was performed using 1x Fast SYBR Green Master Mix (Thermo-Fisher) with gene-specific primers according to the manufacturer’s instructions and performed on a Life Technologies Viia7 Real-time PCR system (Thermo-Fisher). As a reference gene, *spy49_0905* (*gyrA*) was used, as this gene has been previously reported to not have differential expression during growth (55). Nontemplate controls were included to confirm the absence of primer-dimer formation, and all RNA samples were run a minimum of one time as a control to verify absence of high levels of contaminating genomic DNA. All samples were run in triplicate biological and technical replicates on a single plate. Relative gene expression was determined using the 2-ΔΔCT method (56). Data was plotted and statistical significance between expression levels was determined via One-way ANOVA with Tukey’s Multiple Comparisons Post-test with Graph Pad Prism 9.2.0 (GraphPad Software).

### Sample isolation for TMT-LC-MS/MS

To isolate samples for TMT-LC-MS/MS, the following procedure was used and adapted from Wilk *et. al*, 2018 (22). Overnight cultures were diluted 1:100 into 50 mL CDM and grown at 37°C until OD_600_ ∼0.05-0.10, at which time 100 nM SHP peptide was added to required cultures and cells were placed back at 37°C. When cells reached late exponential/early stationary phase (OD_600_ ∼0.70-1.0), cultures were transferred to 15 mL conical tubes and centrifuged at 2700×*g* for 15 min at 4°C. After centrifugation, supernatants were separated from cell pellets into new 15 mL conical tubes.

Supernatants were then filtered through a 0.2 µM pore size filter via syringe (VWR) and Pierce Protease Inhibitor Tablets, EDTA-free (Thermo-Fisher, A32965) were added to 30 mL aliquots of filtered supernatants. Supernatants were placed on ice and subsequently concentrated using Amicon Ultra-4 Centrifugal Filter Units with a 10 kDA MW cutoff (Millipore-Sigma) by centrifugation continually at 2700×*g* for 15 min at 4°C until samples reached 1 ml volume. After samples were concentrated to 1 mL, samples were diluted three times with 5 mL ice-cold PBS and concentrated back down to 1 mL each time. After final concentration to 1 mL with ice-cold PBS, samples were collected, and further concentrated to 250 µL with Amicon Ultra-0.5 Centrifugal Filter Units with 10 kDA MW cutoff (Millipore-Sigma). Final concentrated supernatants were stored at −70°C until proceeding to further sample prep for TMT-LC/MS/MS.

Pellets from isolated samples were washed with 5 mL of ice-cold PBS and centrifuged at 2700×*g* for 15 min at 4°C. Supernatants were discarded and samples were washed twice in 5 mL of chilled TE buffer (50 mM Tris-HCl, pH 8.0; 1 mM EDTA). After the second wash, pellets were resuspended in 1 mL of ice-cold Enzyme Buffer (50 mM Tris-HCl, pH 8.0; 1 mM EDTA; 20% (w/v) sucrose; Pierce Protease Inhibitor Tablets, EDTA-free) with 100 units of Mutanolysin (Sigma-Aldrich, M9901. Resuspended pellets in enzyme mix were transferred to sterile microcentrifuge tubes and incubated at 37°C for 18 hours with shaking (200 rpm). After digestion with mutanolysin, tubes were centrifuged at 21,130×*g* for 5 min at room temperature. Supernatants (solubilized cell wall proteins) were collected and stored at −70°C until proceeding to further sample prep.

After obtaining secreted and solubilized cell wall proteins, total protein concentrations of each sample were measured using the Bio-Rad DC Protein Assay (Bio-Rad, 5000112) according to the manufacturer’s instructions. 100 µg of each protein sample was then submitted to Mass Spectrometry Core in Research Resources Center of University of Illinois at Chicago.

### Sample preparation and digestion for TMT-LC-MS/MS

Filter-aided sample preparation (FASP) was performed with samples submitted to the Mass Spectrometry Core. In brief, protein samples were dissolved by 8 M urea in 0.1 M Tris-HCl, pH 8.5 and then filtered with a 0.22 μM membrane (Millipore). The flow-through was collected and transferred into a 1.5 mL MicroconYM-30 centrifugal unit (Millipore). Protein reduction, alkylation and tryptic digestion were performed step by step in the centrifugal unit. After overnight digestion at 37°C, the peptides were eluted twice with 50 μL 0.1% formic acid (FA). The concentration of proteins and peptides collected in each step was measured using a Nanodrop ONE (Thermo-Fisher). The digested peptides were then desalted, dried, and stored at −80 °C until further use.

### Peptide labeling by TMT, sample fractionation, and TMT-LC-MS/MS data processing and analysis

For isobaric labeling, peptides per sample for 6-plex TMT were labeled using TMTsixplex Isobaric Mass Tagging Kit (Thermo-Fisher, #90064) according to the manufacturers’ instructions. After labeling, excess reagents and detergents were removed by reversed phase solid phase extraction (Oasis HLB C18 SPE, Waters). The samples were dried and resulting pellets stored in − 80 °C.

Sample were then thawed and further fractionated by high pH reversed-phase chromatography using an Agilent 1260 HPLC (Agilent) and a Waters XBridge BEH C18 column (130Å, 3.5 μm, 4.6 x 150 mm). 90 fractions were collected and combined into 10 fractions, followed by desalting using Nestgroup C18 tips (Southborough, MA). Fractionated peptides were dried and redissolved in 0.1% FA for LC−MS/MS analysis.

For LC-MS/MS, fractions were run on Thermo Fisher Orbitrap Velos Pro coupled with Agilent NanoLC system (Agilent) over a 60 min gradient on an Aglient Zorbax 300SB-C18 LC column (15 cm × 75 μm ID, Aglient). Samples were run via a 60 min linear gradient (0–35% acetonitrile with 0.1% FA) and data were acquired in a data dependent manner, in which MS/MS fragmentation was performed on the top 12 intense peaks of every full MS scan. RAW files were converted into .mgf files using MSConvert (57). A database search was carried out using Mascot Server (v2.6, Matrix Science) and Mascot Daemon Toolbox (Matrix Science) for peptide matches and protein searches against the *Streptococcus pyogenes* NZ131 genomic database UniProtKB (Proteome ID: UP000001039). For TMT reporter ion detection, the fragment mass tolerance was set to 20 ppm. Fixed modifications were defined as carbamidomethylation (57.02146) on cysteine and TMT modification (229.1629) on N-terminus and lysine. Miscleavage was set to 2. Search results from 18 runs were imported into Scaffold (Proteome Software) for quantitative analysis.

Peptide identifications were accepted if they could be established at 1% FDR probability as specified by the Peptide Prophet algorithm (58) and as protein identification probability with 99% or greater confidence as determined by Scaffold. Scaffold performed quantitative analysis including extract of the TMT reporter ion intensities from the MS/MS, correction of isotope contamination, and normalization. As equal amounts of total protein were loaded across all TMT channels, reporter ion intensities in each of the channels were normalized by the sum of all reported ion intensities of the corresponding channel. Normalized reporter ion intensities were used for relative protein abundance calculations. Comparisons between groups were made by use of multiple-sample ANOVA test in MaxQuant 1.61 software (59). Statistical significance was set at adjusted q value < 0.05.

Upon receiving the final metadata from the Mass Spectrometry Core in Research Resources Center of University of Illinois at Chicago, the data was submitted to UIC Research Informatics Core. Differential protein expression analysis was then performed using limma package (60). A 1-way ANOVA was used to perform multigroup analysis, as well as pairwise comparisons. P-values were adjusted for multiple testing using the false discovery rate (FDR) correction of Benjamini and Hochberg (61). Protein level differences were considered statistically significant if q values < 0.05. Venn diagrams illustrating sample analysis were created using the BioVenn online platform (54).

TMT-LC/MS/MS data have been deposited to the ProteomeXchange Consortium (http://proteomecentral.proteomexchange.org) via the PRIDE partner repository (62) with the dataset identifier PXD033703 and DOI 10.6019/PXD033703.

### Growth curves of strains

Overnight cultures were diluted 1:100 into 6 mL of prewarmed CDM and grown at 37°C, with OD_600_ observed every 45 min to 1 hour with a GENESYS 30 Vis spectrophotometer. When strains reached OD_600_ ∼0.05-0.1, 100 nM SHP peptide was added to required tubes. Data was plotted and doubling times were calculated by taking OD_600_s ∼0.015 – 0.9 and performing a Nonlinear regression analysis with Exponential growth equation fitting with Graph Pad Prism 9.2.0 (GraphPad Software). Statistical significance between doubling times were determined by One-way ANOVA with Šidák’s or Dunnett’s Multiple Comparisons Post-test with Graph Pad Prism 9.2.0.

### Luciferase assays

Overnight cultures were diluted 1:100 into prewarmed CDM with either 1% glucose or 1% mannose. When cells reached OD_600_ ∼0.05-0.1, 100 nM SHP peptide was added if required. At each time point, OD_600_ measurements were taken with a GENESYS 30 Vis spectrophotometer and luciferase measurements were conducted by removing 100 µL aliquots from each strain and condition and transferred to an opaque 96-well plate. Samples were exposed to decyl aldehyde (Sigma-Aldrich) fumes for one minute and counts per second (CPS) were measured using a Veritas microplate luminometer (Turner Biosystems). Relative light units (RLU) were calculated by normalizing CPS to OD_600_. Data from experiments was plotted and analyzed using Graph Pad Prism 9.2.0.

### Western blotting

Samples were grown as detailed in *Isolation of RNA from bacterial cultures for qRT-PCR and RNA-seq* section, with the following exceptions. To overnight growth of cultures, antibiotics were added if required to select for reporter plasmid propagation. The next day, cells were grown in 6-10 mL CDM, and when cells reached OD_600_ ∼0.05-0.1, 100-120 nM SHP peptide was added if required. Cultures were then grown until OD_600_ reached ∼0.7-1.0. At this time, cultures were harvested and centrifuged at 2700×*g* for 10 min. Supernatants were discarded and cells were resuspended in 250-500 µL Western Blot Lysis Buffer (20 mM Tris-HCl, pH 7.5; 200 mM NaCl), transferred to screwcap tubes with ∼150-250 mg 0.1 mM diameter zirconia/silica beads (BioSpec Products), and subsequently lysed for 10 min in a Mini-Beadbeater-16 (BioSpec Products). After lysis, cells were spun down at 21,130×*g* for 1 min. Supernatant was harvested and transferred to 1.5 mL Eppendorf tubes on ice. Samples were subsequently measured for total protein concentration using the Bio-Rad DC Protein Assay (Bio-Rad).

After determining protein concentration, samples were diluted 1:1 with 2X Laemmli Sample Buffer (120 mM Tris-HCl, pH 6.8; 4% SDS, 20% Glycerol, trace bromophenol blue, 1.43 M β-mercaptoethanol) and boiled for 10 min at 100°C. 15 µg of each sample was then loaded on a 12% SDS-PAGE gel, along with 5 µL of Precision Plus Protein Dual Color Protein Standard (Bio-Rad, #1610374). Loaded SDS-PAGE gels were run with 1X SDS-PAGE Running Buffer (25 mM Tris-HCl, pH 8.3; 192 mM glycine; 0.1% SDS) at 150 V for 1.5 hours on a Mini-PROTEAN Tetra Vertical Electrophoresis Cell (Bio-Rad) with a PowerPac Basic Power Supply (Bio-Rad). Proteins separated by SDS-PAGE were transferred to nitrocellulose membrane (0.45 µm, Thermo-Fisher, #88018) at 350 mA for 1.5 hours using a Mini Trans-Blot Module (Bio-Rad) hooked up to the PowerPac Basic Power Supply. Membranes were subsequently washed twice with TBST (50 mM Tris-HCl, pH 7.5; 150 mM NaCl, 0.05% Tween-20) and blocked with TBST + 5% BSA (bovine serum albumin, A2153) for 30 min. Blots were then rinsed briefly twice with TBST and incubated with 1:4000 rabbit anti-GFP-Tag pAb (ABClonal, #AE011) in TBST + 5% BSA overnight at 4°C. The next morning, blots were washed three times for 10 min with 10 mL TBST, and then incubated with 1:100,000 goat anti-rabbit IgG (H+L) secondary antibody conjugated to HRP (Thermo-Fisher, #31460) for 1 hour. Blots were washed again with 10 mL TBST three times for 10 min, and subsequently incubated with SuperSignal West Femto Maximum Sensitivity Substrate (Thermo-Fisher, #34094) according to the manufacturer’s instructions. Blots were then imaged using a ProteinSimple FluorChem R Imaging System (ProteinSimple, Biotechne) on the chemiluminescence setting for 1-5 min. After imaging, blots were briefly washed twice with 10 mL TBST and stained for protein loading with India ink (Thermo-Fisher, R21518) in TBST. India ink stained blots were imaged using a G:BOX Chemi XRQ gel doc system (Syngene) and GeneSys (ver. 1.5.6.0) software on the white light setting for 175 msec.

### Preparation of radiolabeled (p)ppGpp standards from *E. coli*

*E. coli* (p)ppGpp standards were prepared as follows, and the procedure was adapted from Kaczmierczak *et. al*, 2009 (45). Strain TX2737 (Table S1) was streaked to single colony isolation from frozen glycerol stocks at −70°C on LB + Amp plates, and grown overnight at 37°C. The next day 5-6 colonies were suspended in 200 µL of MOPS-CDM 150 µCi/mL [^32^P] orthophosphate (Perkin Elmer, NEX053H001MC) and with or without 100 mM IPTG. Cells were incubated at 37°C for 30 min, and at this time, 25 µL culture aliquots were harvested and placed into tubes containing 25 µL ice-cold 13 M formic acid. Following this, three consecutive freeze thaw cycles alternating cells on dry ice vs. incubation at 37°C were performed. After the final freeze-thaw, cells were spun down at 8,000×*g* for 5 min. After centrifugation, supernatant was carefully removed without resuspending cellular debris and transferred to new tubes. Standards were diluted 1:1 with 13 M formic acid and stored at −20°C for up to 2 weeks.

### Non-uniform labelling of (p)ppGpp from *S. pyogenes* and TLC

*S. pyogenes* was grown and radiolabeled nucleotides were extracted as follows, adapting the procedure from Kaczmierczak *et. al,* 2009 (45). Pre-stored glycerol stocks of bacterial strains at −70°C were inoculated into THY broth and incubated overnight at 37°C. The next morning, overnight cultures were centrifuged at 2700×*g* for 10 min and washed twice with 1 mL of prewarmed CDM. Strains were diluted 1:200 into 6 mL of fresh prewarmed CDM and grown at 37°C, until they reached OD_600_ ∼0.05-1.0, and then 100 nM SHP was added to required cultures. Cells were grown for another 1.5-2 hrs (when wild-type reached OD_600_ ∼0.3-0.5) and cells were centrifuged again at 2700×*g* for 10 min. After centrifugation supernatants were discarded, and cells were resuspended in 1 mL of MOPS-CDM. Cultures were centrifuged again at 2700×*g* for 10 min, supernatants were discarded, and cultures were resuspended in 6 mL MOPS-CDM or MOPS-CDM containing 100 nM SHP peptide. OD_600_ was measured using a spectrophotometer and cells were adjusted to the same density (around OD_600_ ∼0.05) in 1 mL volumes. 92.5 µL of each culture were transferred to new Eppendorf tubes, and 7.5 µL of 2 µCi/µL (final concentration: 150 µCi/mL) [^32^P] orthophosphate was added to each culture. Cultures were incubated at 37°C for 30 min (according to the maximal induction of (p)ppGpp seen by Steiner and Malke, 2000 (28)), and then cultures were harvested and transferred to Eppendorf tubes containing 100 µL ice-cold 13 M formic acid. Following this, three consecutive freeze thaw cycles alternating cells on dry ice vs. incubation at 37°C were performed. After the final freeze-thaw, cells were spun down at 8,000×*g* for 5 min. After centrifugation, supernatant was carefully removed without resuspending cellular debris and transferred to new tubes that were kept on ice or at −20°C until running on TLC.

For TLC separation, a PEI-cellulose TLC plate (Millipore-Sigma, #Z122882) pre-run in distilled H2O and dried was obtained. To the TLC plate, 2-3 µL of each sample and *E. coli* standards were added. TLC was developed with 1.5 M KH_2_PO_4_, pH 3.4; removed from the chromatography chamber and allowed to dry overnight. The next morning, the plate was exposed to a storage phosphor screen (Molecular Dynamics) for at least 4 hours. The storage phosphor screen was subsequently imaged using an Amersham Typhoon scanner (GE) set to Phosphor Imaging scanning mode with the following settings: 4000 sensitivity, 200 µM pxl size. Resulting images were contrast adjusted using ImageJ (63).

## ACKNOWLEDGEMENTS

The authors would like to acknowledge the following research cores for their help in data acquisition. RNA-seq library preparation and data collection was supported by the Northwestern University NUSeq Core Facility; TMT-LC-MS/MS data acquisition and partial analysis was performed by the Mass Spectrometry Core in the Research Resources Center of University of Illinois at Chicago; and additional bioinformatics analysis of TMT-LC-MS/MS was performed by the UIC Research Informatics Core, supported in part by NCATS through Grant UL1TR002003. We also thank M.E. Winkler and H.T. Tsui for the kind gift of the *E. coli* strain TX2737. Additionally, we thank the members of the Federle lab for helpful discussions and proofreading of the manuscript.

## FUNDING AND ADDITIONAL INFORMATION

This study was supported by the National Institutes of Health (R01-AI091779 to M.J.F. and 5K12 GM139186/NIGMS/NIH to B.E.R.) and the DoD’s Science, Mathematics and Research for Transformation (SMART) Scholarship for Service Program to C.M.A.

